# Alternative silencing states of Transposable Elements in Arabidopsis

**DOI:** 10.1101/2024.03.16.585326

**Authors:** Valentin Hure, Florence Piron-Prunier, Tamara Yehouessi, Clémentine Vitte, Aleksandra E. Kornienko, Gabrielle Adam, Magnus Nordborg, Angélique Déléris

**Author notes:** Contributed equally (Valentin Hure, Florence Piron-Prunier, Tamara Yehouessi).

## Abstract

The DNA methylation/H3K9me2 and Polycomb-group proteins (PcG)-H3K27me3 pathways have long been considered mutually exclusive and specific to TEs and genes, respectively. However, H3K27me3 can be recruited to many TEs in the absence of DNA methylation machinery and sometimes also co-occur with DNA methylation. In this study, we show that TEs can also be solely targeted by H3K27me3 in wild-type Arabidopsis plants. These H3K27me3-marked TEs not only comprise degenerate relics but also seemingly intact copies that display the epigenetic features of responsive PcG target genes as well as an active H3K27me3 regulation. We also show that H3K27me3 can be deposited on newly inserted transgenic TE sequences in a TE-specific manner indicating that silencing is determined in *cis*. Finally, comparison of Arabidopsis natural accessions reveals the existence of a category of TEs - which we refer to as “*bifrons*” - that are marked by DNA methylation or H3K27me3 depending on the ecotype. This variation can be linked to intrinsic TE features and to *trans*- acting factors, and reveals a change in epigenetic status across TE lifespan. Our study sheds light on an alternative mode of TE silencing associated with H3K27me3 instead of DNA methylation in flowering plants. It also suggests dynamic switching between the two epigenetic marks at the species level, a new paradigm that might extend to other multicellular eukaryotes.

## INTRODUCTION

Transposable Elements (TEs) are repeated sequences that can potentially move and multiply in genomes. Transposition is typically deleterious, and is tightly controlled by host organisms. On the other hand, TE mobilization can be an important factor in evolution and adaptation, in particular by providing nearby genes with genetic or epigenetic regulatory modules that can impact transcriptional programs^1^. Amplification of a TE typically leads to the production of groups of copies that are identical in sequence upon insertion, thus producing so-called ‘TE families’. Over time, TE insertions accumulate SNPs and indels, leading to the generation of mutated and truncated copies. These are often non-autonomous, either due to their lack of functional protein-coding genes, structural regions necessary for their life cycle, or both. In the first case, copies can nevertheless be mobilized using proteins produced in *trans* by copies of the same family. In the absence of new insertions, eventually only highly degenerate relics with no potential for mobility will remain^2^; they will also disappear unless selectively maintained for their *cis* (or *trans*) regulatory impact^3^.

A hallmark of TE silencing in many organisms is high DNA methylation at cytosines (5-methylcytosines). In plants, TE methylation can be found in three sequence contexts (CG, CHG and CHH, where H can be any base except G), and is coupled with histone H3K9 dimethylation (H3K9me2). The combination of these two epigenetic marks prevents TE expression and transposition, thus guaranteeing genome integrity^4,5^. DNA methylation is established at TEs upon their insertion in the genome by the RNA-directed DNA methylation pathway: TE-derived small RNAs (or small RNAs pre-existing in the nucleus) guide methyltransferase DRM2 to homologous TE sequences to establish DNA methylation in all three sequences cytosine contexts^6–8^. Four different pathways amplify the established DNA methylation patterns and maintain them stably over time. CG, CHG and CHH DNA methylation patterns are typically maintained by MET1^9^, CMT3^10^, and CMT2 and RdDM^11^, respectively. While very stable across generations, DNA methylation can be removed by DNA glycosylases of the DEMETER-like family (DML): these constitutively prune the DNA methylation patterns^12^, prevent spreading outside of primary targeted sequences^13^ or actively participate to gene control, either at specific stages of development or by preventing stress-responsive genes from being locked in a constitutively silent state.

DNA methylation and H3K9me2 patterns are known to be stable throughout development and generations, and are hallmarks of the so-called ‘constitutive heterochromatin’. On the other hand, H3K27me3 is associated with facultative heterochromatin and is a hallmark of transcriptional repression associated with protein- and miRNA-coding genes involved in development, reproduction or metabolic and stress responses^14–17^. Presence of H3K27me3 is tightly regulated by a balance between active addition by Polycomb Repressive Complex 2 (PRC2)^18^ and active removal by histone REF6-ELF6-JMJ13/30/32 demethylases of the KDM4/JMJD2 family. H3K27me3 deposition establishes cell identity and silencing of stress-responsive/metabolic genes and its removal can allow gene activation^19^, contribute to shape the spatial and temporal patterns of H3K27me3, reset the mark during reproduction^20,21^, and maintain H3K27me1 patterns^22^.

Because of their different targets and contrasted properties, PcG-H3K27me3 and DNA methylation have been generally considered as mutually exclusive systems, specialized for the transcriptional silencing of genes and TEs, respectively. However, this dichotomous model has been challenged in the past decade. In both plants and mammals, targeting of PcG to TEs was initially evidenced by gain of H3K27me3 at many TE sequences upon their loss of DNA methylation^23–29^ in mutants unable to maintain DNA and H3K9 methylation, as well as in specific cell types where TEs are naturally demethylated^25,30^. This implies that PcG could serve as a back-up silencing system for hypomethylated TEs (compensation), and functional redundancy of the two systems has been indeed revealed at some TEs in both mammals and plants^3,31,32^. These observations also have two mechanistic implications: that PcG can be recruited to TEs, and that DNA methylation can exclude H3K27me3 deposition. Yet, in some instances, the two marks can co-occur^33,34^ and cooperate in restricting TE activation upon biotic stress^3^. More recently, it was also reported that PcG, instead of the DNA methylation machinery, targets and represses TE in wild type unicellular eukaryotes and plants from early lineages, thus illustrating an ancient role of PcG in regulating TE silencing^35,36^.

TE targeting by PcG has been well described in DNA methylation mutants of *A. thaliana*, but whether this can also occur in the presence of an active DNA methylation machinery, replacing DNA methylation, remains unclear. Moreover, the modes of PcG recruitment at TEs remain to be fully elucidated: while some TEs seem to be marked by H3K27me3 as the result of neighboring gene spreading^3^, observation of H3K27me3 mark at TEs located away from any H3K27me3 marked gene indicates possible specific *cis*- recruitment, but this has not been formally tested. Besides, recruitment of PcG to TEs in the absence of DNA methylation points to a possible *de novo* recruitment on newly inserted TE sequences but this is unknown and needs to be investigated.

In this study, we report that many transposable elements (>4 000) are covered by the H3K27me3 histone mark in the *A. thaliana* genome in WT plants. These elements comprise not only short TE relics, but also intact copies, which are expected to be targets of DNA methylation. We show that these long copies present similarities to classical PcG genic targets: they display the chromatin hallmarks of genes responsive to environmental or developmental cues and their H3K27me3 patterns are actively regulated by histone H3K27 demethylases. Using transgenic constructs, we demonstrate that newly inserted TE sequences are able to recruit H3K27me3 *de novo,* with patterns specific to each copy, indicating that recruitment of H3K27me3 relies on intrinsic features of the TE copies themselves. Finally, natural variation between Arabidopsis accessions at orthologous TE copies reveal an epigenetic switch between the two silencing modes. Using a GWAS approach, we provide evidence that this epigenetic switch is linked to both *cis*- and *trans*- determinants.

Our results establish an alternative, *cis*-determined epigenetic control of TEs by PcG in wildtype *A. thaliana;* they further provide evidence that a given TE can change its epigenetic silencing state throughout its lifespan. This challenges the dogma that Arabidopsis TE are solely targeted by DNA methylation in nature.

## RESULTS

### Many TEs are marked solely by H3K27me3 in wild-type Arabidopsis plants

When examining H3K27me3 ChIP-seq profiles generated for natural Arabidopsis accessions, we noticed that several TEs showed an absence of DNA methylation and were instead marked by H3K27me3 (**Fig.1A** and **EV1A**). To determine their abundance in the genome, we defined four classes of TEs based on the length coverage of H3K27me3 marks and CG methylation, using these datasets and previously published methylomes^37^ (**Fig.1B** and **EV1B**). The class which consists of TEs marked by H3K27me3 (“Class 3”) represents 15% of the total TE number but only 8.4% of TE bases. These percentages indicate a bias in the size of the TEs targeted H3K27me3 (**Fig.1B** versus **EV1C**). Accordingly, TE with H3K27me3 tend to be more present within short TE categories (particularly those with size below 1kb), while long TEs (with size over 3.5kb) are generally associated with DNA methylation (**Fig.1C**)—although a small fraction of long TEs (>3.5 kb) are marked by H3K27me3 alone (**Fig.1C** and **EV1D**).

**Figure 1.**
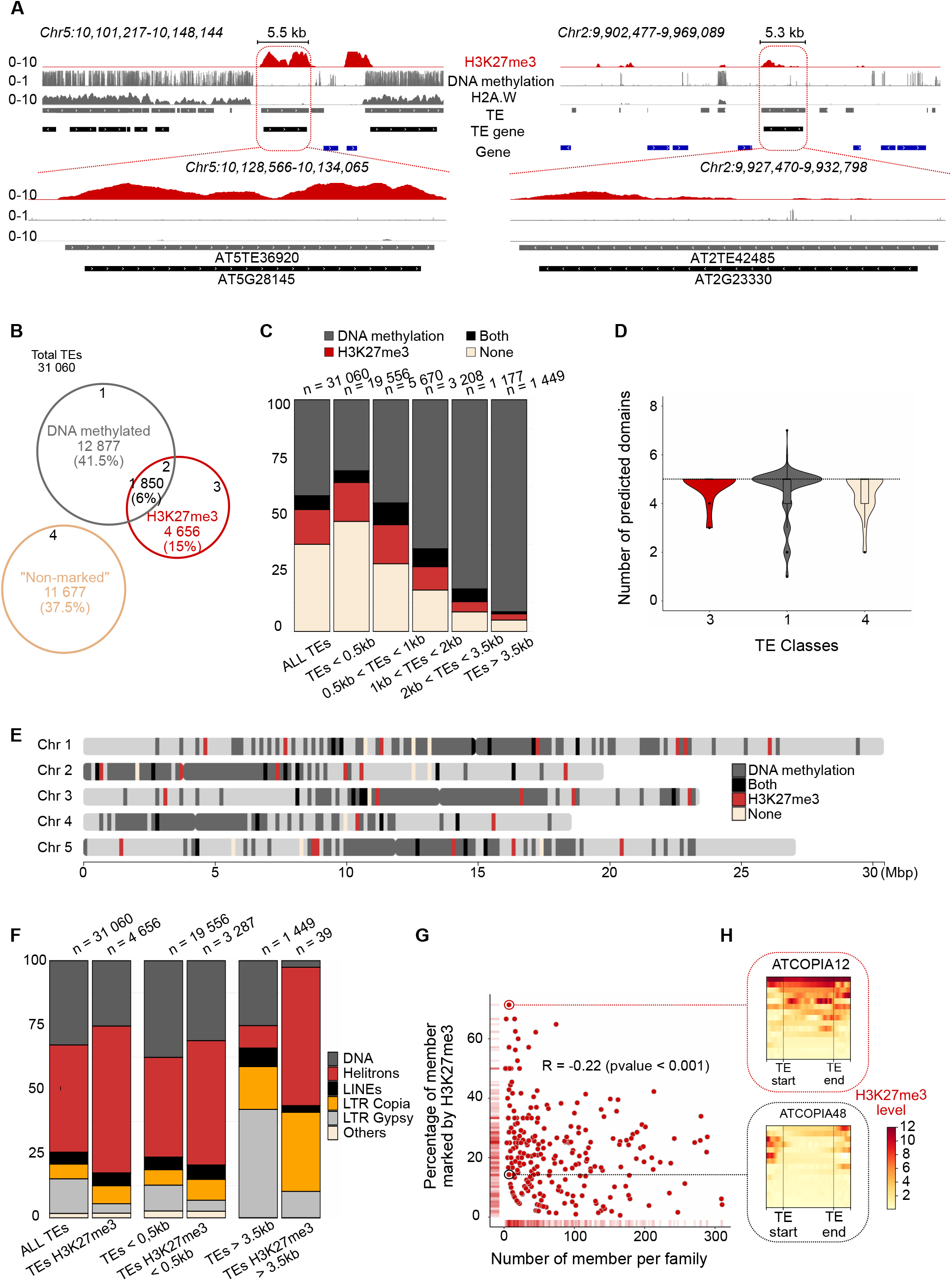
Many TEs are marked by H3K27me3 in wild type Arabidopsis **(A)** Representative genome browser view of H3K27me3 (red), DNA methylation (grey) and constitutivehetero- chromatin histone variant H2A.W (grey) levels in wild-type (WT) for two TEs marked by H3K27me3 in WT. Grey, black and blue bars represent TEs, TE-genes and Protein Coding Genes annotations, respectively. **(B)** Venn diagram shows the classes of TEs based on their epigenetic modifications. TE methylation levels for CHG and CHH in each of these classes are shown in Fig.EV1B. **(C)** Stacked bar plot shows distribution of TE classes according to TE length. **(D)** Violin plot shows number of predicted domains for LTR Copia among TE>3.5kb using DANTE (Neumann et al, 2019). **(E)** Schematic chromosomes show distribution of TE>3.5kb along genome. **(F)** Stacked bar plot shows distribution of TE super-families according to TE length among all TEs or H3K27me3 marked TEs. **(G)** Plot shows correlation between percentage of members marked by H3K27me3 and number of TE copies per subfamily. Each dot is a TE family (< 300 members). **(H)** Heatmaps based on H3K27me3 levels along each TE copy within one given TE family. **Top panel** : *ATCOPIA12* and **bottom panel** : *ATCOPIA48*.

Interestingly, inspection of their predicted protein-coding domains showed that these TEs contain a number of functional domains similar to that of mobile TEs (**Fig.1D**). Thus, TEs which are expected targets of DNA methylation can also be marked with H3K27me3.

Interestingly, H3K27me3-marked long TEs (>3.5 kb) are dispersed throughout the euchromatic arms and pericentromeric regions (**Fig.1A** and **Fig.1E**), in contrast to the DNA-methylated ones, which are generally clustered in pericentromeres. This suggests that genomic location is not the major determinant of PcG targeting. Further analysis of H3K27me3-marked long TEs showed that they are enriched in Helitrons for all size categories, and particularly enriched in Helitrons and Copia LTR retrotransposons for long copies (**Fig.1F**). Comparison of epigenomic features at copies from the same family revealed intrafamily epigenetic variation (**Fig.EV1E**), with some TE copies within a family being marked by H3K27me3 while others are not. We further observed a weak but significant anti-correlation between copy number and percent of copies marked with H3K27me3 (**Fig.1G**) suggesting that small TE families are more prone to be marked by H3K27me3 than families with large copy numbers. Nevertheless, small family size cannot fully explain epigenomic preference, as families of similar copy number can exhibit very different H3K27me3 levels (exemplified by *ATCOPIA12* and *ATCOPIA48* families, **Fig.1H**). Thus, while the TE characteristics tested could influence epigenetic status, they are not sufficient to explain it, suggesting that additional intrinsic features of TE copies participate in H3K27me3 recruitment.

### H3K27me3 marks at TEs behave like PcG target genes

H3K27me3 was previously shown to colocalize with other chromatin marks associated with transcriptional repression at protein-coding genes. These include H2A.Z -a histone variant associated with responsive genes and antagonized by DNA methylation^38^- and H2AK121ub, which is deposited by the Polycomb repressive complex PRC1^39^ and confers to genes a transcriptional state responsive to developmental or environmental cues^15^. By reanalyzing publicly available datasets, we observed that 23% and 22% of the H3K27me3-marked TEs also harbor H2A.Z or the H2AK121ub mark, respectively (**Fig.2A**). Interestingly, H3K27me3 long TEs are most often marked by H2A.Z and/ or H2AK121ub (**Fig.2A** and **Fig.2B**). Hence, they behave as PcG genic targets, for they display the epigenetic hallmarks of gene responsiveness (**Fig.2C)**.

**Figure 2.**
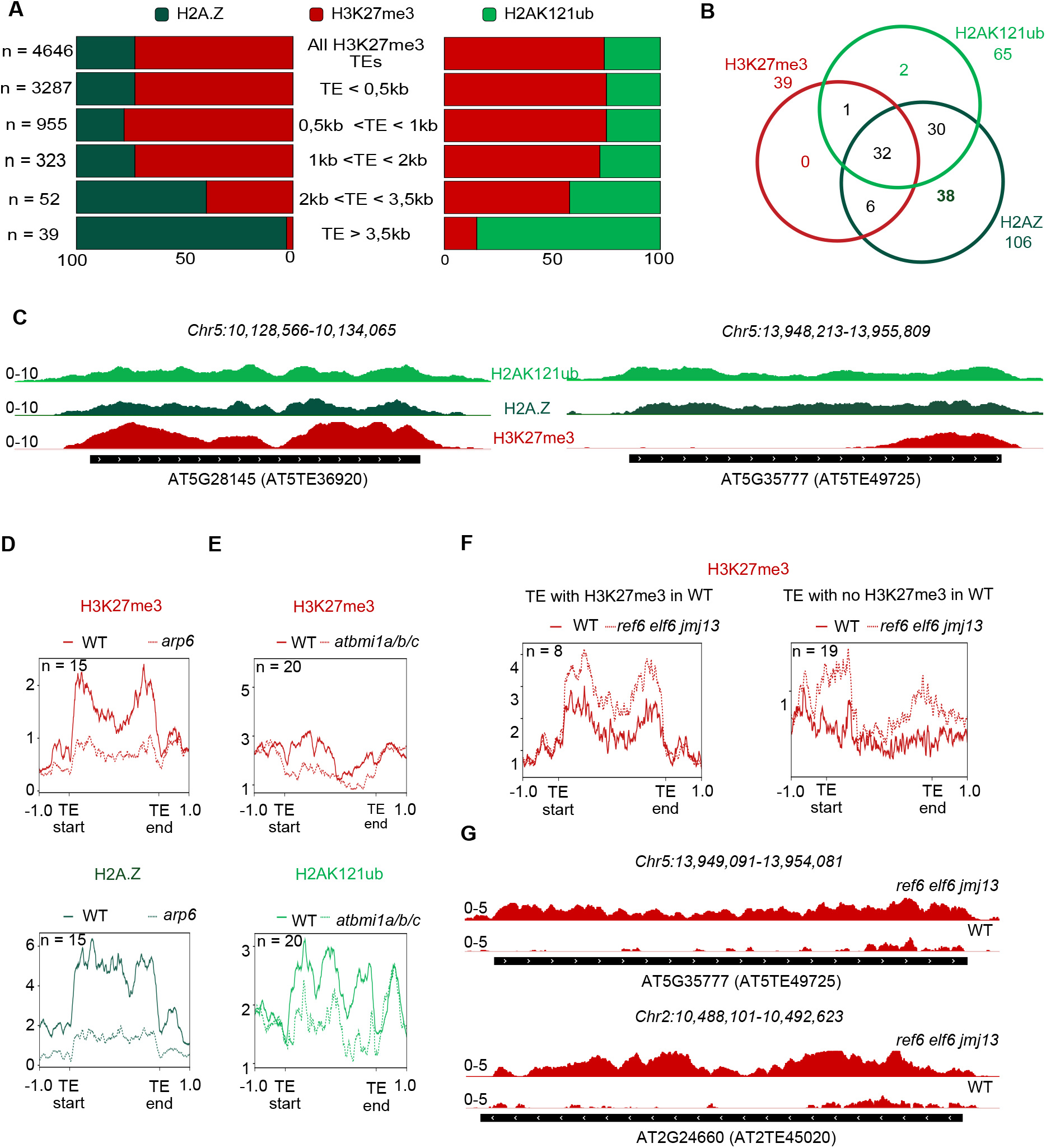
H3K27me3 at TEs can be associated with H2AUb and H2AZ and actively demethylated (A) Stacked bar plot shows distribution of TE marked by, either H2A.Z variant or H2AK121 ubiquitination, among TE marked by H3K27me3 per size range. (**B)** Venn diagram shows presence of H3K27me3, H2A.Z variant and H2AK121ub on TEs>3,5kb. **(C)** Representative genome browser view of 2 TEs from class 3 (marked by H3K27me3) with CHIP-seq data showing H2AK121ub (light green), H2A.Z (dark green), H3K27me3 (red) marks in wild-type (WT). **(D)** Metagenes show level of H3K27me3 (**top panel**) and H2A.Z (**bottom panel**) in both WT and *arp6* mutant defective for H2A.Z incorporation. **(E)** Metagenes show level of H3K27me3 (**top panel**) and H2AK121ub (**bottom panel**) in both WT and *atbmi1a/b/c* mutant defective for H2AK121Ub deposition. **(F)** Meta- genes show level of H3K27me3 in both WT and *ref6 elf6 jmj13* triple H3K27me3 demethylases mutant. **Left panel** represents TEs already marked by H3K27me3 in WT but the mark is increased in *ref6 elf6 jmj13.* **Right panel** represents TEs that are not marked by H3K27me3 in WT but the mark is revealed in *ref6 elf6 jmj13*. **(G)** Representative genome browser view of 2 TEs from class 3 showing H3K27me3 (red) marks in wild-type and *ref6 elf6 jmj13*.

Previous studies showed that, for a subset of genes, recruitment of H3K27me3 is dependent on the presence of H2A.Z^40^ or H2AK121ub^41^. This prompted us to analyze H3K27me3 profiles at TEs in *arp6* mutant impaired for H2A.Z deposition, as well as in *atbmi* PRC1 mutants impaired in H2AUb deposition. At a subset of TEs, we observed a decrease of H3K27me3 in *arp6* **(Fig.2D** and **EV2A)** and *atbmi* plants **(Fig.2E** and **EV2B**), thus revealing similar dependencies for H3K27me3 deposition as for genes. Moreover, given that REF6 was shown to preferentially remove H3K27me3 at H2AK121ub-marked genes^42^, we re-analyzed published data in the triple mutant *ref6 elf6 jmj13* (ELF6 and JMJ13 are known to function redundantly with REF6)^43^. We observed that several long TEs marked by H3K27me3 have higher levels of H3K27me3 in *ref6 elf6 jmj13* in comparison to WT (**Fig.2F left panel** and **Fig.2G**). Thus, the observed profile of H3K27me3 at some TEs is shaped and regulated through active demethylation by JMJ H3K27me3 demethylases (**Fig.2G)**. Interestingly, we also observed a small gain of H3K27me3 in the triple mutant at TEs marked by H2AK121ub but not by H3K27me3 in the WT (55% of cases) (**Fig.2F right panel** and **EV2C**); this shows that the activity of H3K27me3 demethylases can remove H3K27me3 completely so that TEs look like active genes in WT.

These results demonstrate that long (>3.5kb) TEs can harbor chromatin hallmarks of typical PcG genic targets, including active regulation by histone demethylase. These features distinguish them from constitutively heterochromatic TEs.

### H3K27me3 mark is recruited at neo-inserted transgenic TEs in a *cis*-determined manner

Since TEs can recruit PcG in the absence of DNA methylation, we envisioned that PcG targeting could occur *de novo* at newly inserted TE copies, possibly in competition with DNA methylation. To test whether newly inserted TE copies devoid of DNA methylation upon insertion can recruit PcG, we used LTR retrotransposon-based transgenes which can model TE insertion^44^. We further reasoned that such transgenes could enable us to validate the hypothesis raised by Figure 1, which is that recruitment of PcG on TE is instructed by intrinsic features of the TE. For this purpose, we used three *COPIA* LTR retrotransposon candidates that were chosen based on their genomic and epigenomic features. *ATCOPIA12B* (*AT2TE45020*) was identified in **Fig.1H** as part of a family prevalently targeted by PcG in WT and harboring H3K27me3 (**Fig.EV3A top panel**), particularly in *ref6 elf6 jmj13* mutants in which the masking effect of REF6 activity is eliminated (**Fig.2G**). *ATCOPIA21A* (*AT5TE65370*) is an active TE that is DNA methylated in WT but targeted by PcG in *ddm1* with a peak centered on the TE – suggesting that H3K27me3 does not come from nearby gene– (**Fig.EV3A bottom left panel**) and was previously shown to transpose in *ddm1* mutant lines as well as in *ddm1*-derived EpiRILS^45,46^. Finally, *ATCOPIA13A* (*AT2TE23885*) is DNA methylated in WT and targeted by PcG in *ddm1,* with different H3K27me3 patterns compared to *ATCOPIA21* and possibly with a contribution of H3K27me3 spreading from the nearby gene (**Fig.EV3A bottom right panel**).

The three sequences were cloned separately in binary vectors and a SNP was introduced to discriminate transgenes from endogenous sequences (**Fig.3A**). We exploited the SNP to discriminate and quantify H3K27me3 levels on the transgene as compared to the endogenous original sequence. Besides, the H3K27me3 analysis was performed in pools of 15-20 plants to dilute positional effects linked to transgene insertion site, in particular H3K27me3 spreading from nearby regions. Results show consistent recruitment of H3K27me3 *de novo* on the transgenic TEs, although to different extents as quantified using reads counts (**Fig.3B-C** and **Fig.EV3B-D, right panels**). While H3K27me3 spreading from regions adjacent to transgenic copies is possible and could contribute the signal, the reproducible and consistent differences in H3K27me3 patterns between the three TEs indicate that the recruitment of H3K27me3 is specific to each TE (**Fig.3B-C,** and **Fig.EV3B-D, left panels**).

**Figure 3.**
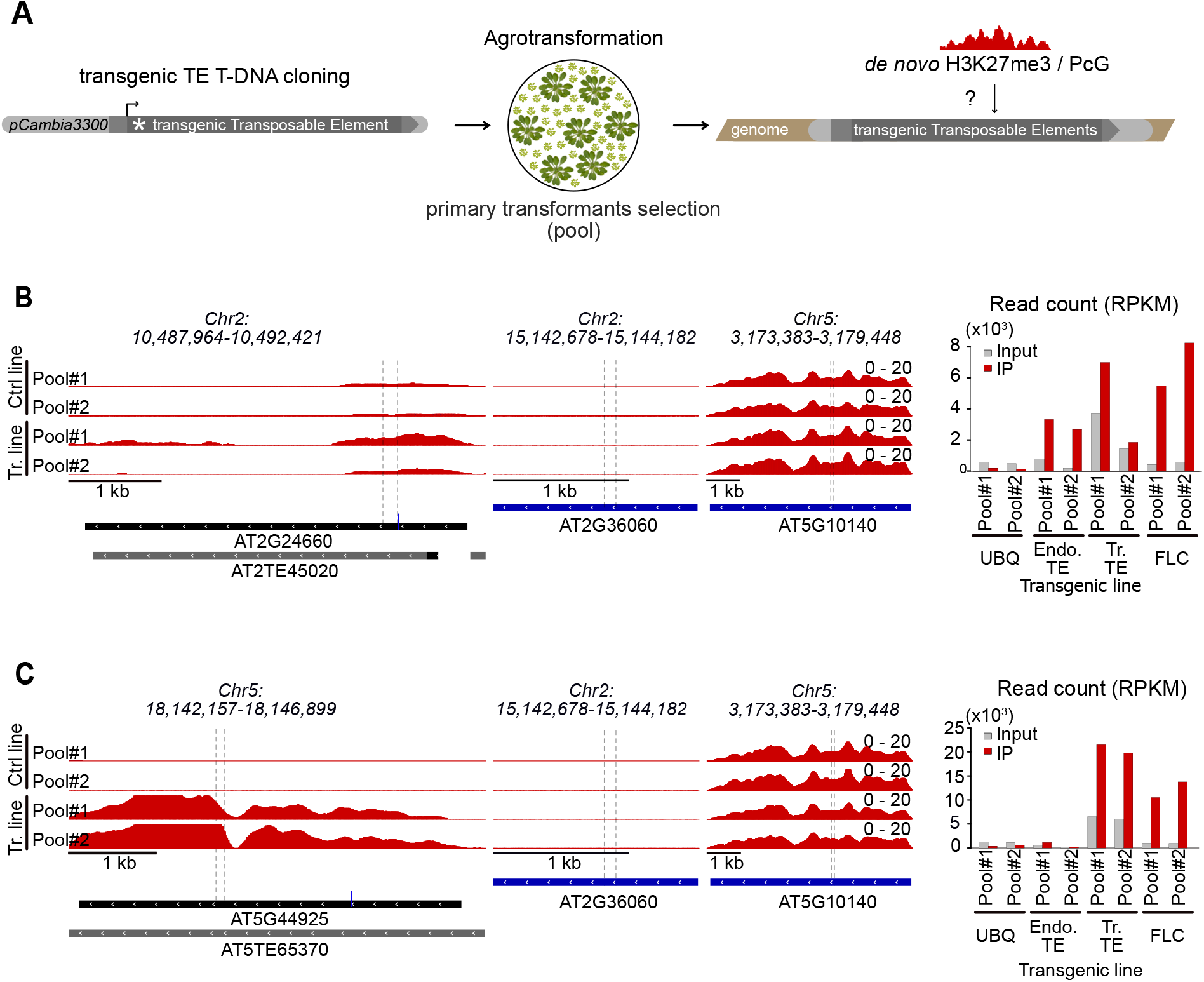
Neoinserted TE sequences can recruit H3K27me3 de novo and in an instructive manner (A) Experimental scheme: individual TE sequences were cloned into a binary vector, introduced in plant genome and studied by H3K27me3 ChIP-seq in large pools of T1(15-20 plants) to dilute position effects. **(B) Left panel:** Representative genome browser view of H3K27me3 average profile (IP-INPUT) at *COPIA12B* sequence (**top panel**) in two independent pools of plants either transformed with unrelated transgene (Ctrl) or with the TE of interest. The H3K27me3 signal in the “Ctrl” tracks reflects the H3K27me3 status at the endogene TE. The H3K27me3 signal in the “Tr.” tracks reflects the H3K27me3 status at the endogene + transgene: higher signal compared to control line indicates recruitment of H3K27me3 on transgenic TEs in addition to the endogene. Variability between transgenic pools reflects the difference in transgenic TE copy number which varies between Pool 1 and 2. **Middle panel**: *UBQ* and *FLC* are shown are negative and positive controls respectively for H3K27me3 recruitment. **Right panel:** Recruitment on the transgene is by quantification of endogenous and transgenic *COPIA12B* sequences immunoprecipitated with H3K27me3 using CHIP-seq reads at the region where a SNP was introduced to discriminate transgene and endogene. Reads overlapping with the SNP at *COPIA12B* locus were extracted, counted and normalized by total read number. Recruitment on the transgene is validated by enrichment observed in IP as compared to an Input fraction. At control regions (*UBQ* and *FLC*), reads were extracted at positions *Chr5:3,178,750* (1^st^ intron) and *Chr2:15,143,214* respectively (dashed lines). Copy number variation between transgenic pools can be visualized by the input DNA fraction (grey bar) / In the non-transgenic lines, variation of input reflects difference in sequencing coverage at the SNP. **(C) Left**, **middle** and **right panels** are as in (B) except for *COPIA21* sequence.

### Switching between DNA methylation and Polycomb at TEs at the population level

The observation of relatively intact H3K27me3-marked TEs, which have gene-like chromatin features, raises the question of whether this epigenetic state at TE is conserved across accessions given that H327me3 genic targets are conserved across Arabidopsis species^16^.

To study this, we first compared Col-0 to KBS-Mac-74 - an accession that was recently profiled for H3K27me3, H3K9me2 and DNA methylation^36^ and for which high-quality ONT-based genome assembly was available^47^. Using these data, eight clusters of TEs were defined based on H3K27me3 and H3K9me2 (used as a proxy for DNA methylation) profiles (**Fig.4A, 4B** and **4C**). In the two largest clusters (1 and 2) epigenomic features remain unchanged; clusters 3 and 4 display epigenetic variations which are likely contributed by genetic diversity between both ecotypes, (*i.e.,* TEs that are present in Col-0 and absent in KBS-Ma-74, change in copy number and/or TE size); clusters 5 and 6 show variation of one mark but not the other. Strikingly, the last 2 clusters (7 and 8) (67 and 50 TEs, respectively) correspond to TEs with a change in epigenetic status between the two compared accessions **(Fig.4C**). Thus, we have unveiled a sub-population of TEs with an epigenetic status that is variable on the species-level (**Fig.4D** and **Fig.EV4A** for additional examples): we coin these TE as “*Bifrons*” which literally means “two-faced” in Latin. This phenomenon observed at orthologous TE insertions indicates a change in the epigenetic state – from standard TE-like DNA methylation to H3K27me3, which we refer to as an “epigenetic switch”. Interestingly, the epigenetic switch can be specific to the TE locus, with no observable epigenetic or transcriptional change at nearby regions (**Fig.EV4B**).

**Figure 4.**
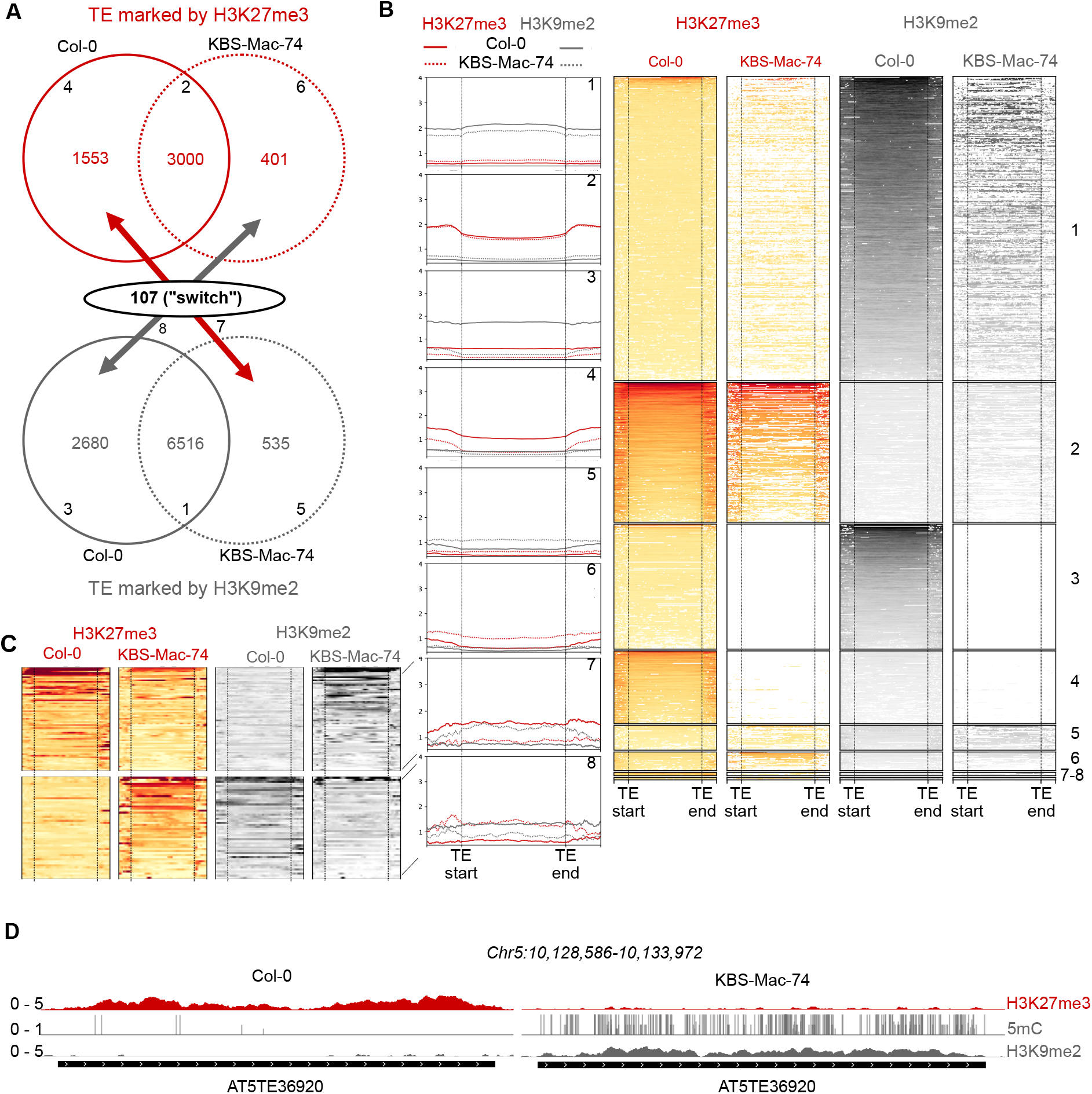
Switch between silencing epigenetic marks at TEs visualized by inter- accessions comparisons (A) Venn diagram of TEs marked by H3K27me3, at the top (numbers in red), or by DNA methylation (H3K9me2 used as proxy) at the bottom (numbers in grey), in 2 different ecotypes, Col-0 and KBS-Mac-74. Intersection between both Venn diagrams show TEs (107) that switch from H3K27me3 in one ecotype to H3K9me2 in the other. **(B)** Metaplots and heatmaps corresponding to the different Clusters 1-8 shown in (B) based on H3K27me3 and H3K9me2 values for TEs in both ecotypes (white bars represent missing coverage). **(C)** Zoom-in on heatmaps for clusters 7 and 8 which correspond to TEs that undergo the epigenetic switch between the two ecotypes. **(D)** Representative genome browser view of H3K27me3 and DNA methylation levels at an orthologous TE copy in two ecotypes, Col-0 and KBS-Mac-74, respectively (red : H3K27me3, dark grey: CG methylation, grey: CHG methylation, light grey: CHH methylation, black bar: TE annotation).

Other pairwise comparisons such as Col-0 versus Fly-2 or KBS-Mac-74 versus Fly-2, showed a similar number of TEs that differ in epigenetic state (**Fig.EV4E**). Based on this, we conclude that the *Bifrons* are a rather small sub-population of TEs, but they likely occur in all natural accessions of Arabidopsis.

### H3K27me3 deposition at some TEs instead of DNA methylation is genetically linked to TE *cis*-determinants as well as *trans* factors

Next, we wanted to identify possible genetic determinants of this epigenetic variation. For this purpose, we exploited publicly available DNA methylation data for hundreds of Arabidopsis natural accessions and performed a Genome-Wide Association Study (GWAS). First, we took the opportunity that H3K27me3 and H3K9me2/DNA methylation data were recently profiled in 12 ecotypes^37^ to test whether the presence of DNA methylation/H3K9me2 is generally correlated with the absence of H3K27me3 and vice-versa. To do this, for six individual ecotypes, we plotted H3K27me3 levels against these of H3K9me2 for all the TEs that have either one of the marks, and indeed observed an anticorrelation between H3K27me3 and H3K9me2/DNA methylation (**Fig.EV5A**). This anticorrelation allows us to extrapolate a H3K27me3 status from the absence of DNA methylation at TEs in nearly 500 ecotypes for which DNA methylation but not H3K27me3 data was available. Thus, we could perform GWAS at a given TE, using presence/absence of DNA methylation as a molecular phenotype. We focused our analysis on two individual TEs of the *COPIA12* family that display genetic and epigenetic variation. *ATCOPIA12D* (*AT5TE36920*) insertion (shown in **Fig.4D**) is present in all the assembled accessions and is always DNA methylated except in Col-0 where it is marked by H3K27me3. *ATCOPIA12B* (*AT2TE45020*) insertion is either present or absent and there can be multiple insertions of the corresponding sequence which can be found either in a DNA methylated state and/or H3K27me3 state. DNA methylation level was estimated for CG, CHG, CHH respectively in each ecotype for the two TEs (**Fig.5A**). At *ATCOPIA12D* (*AT5TE36920*), a clearly bimodal distribution of DNA methylation was observed: most of the accessions are DNA methylated in all three contexts but a small fraction of ecotypes has methylation levels close to zero (**Fig.5A**), likely with H3K27me3 instead as suggested by our previous results (**Fig.4**). GWAS analysis showed a distinctive peak *in cis* located on the TE itself (**Fig.5B**). In addition, other smaller peaks could be observed close to genes known to be important players of the DNA or H3K9 methylation pathway, such as *RDR2, CMT2 and SUVH5* (**Fig.EV5B**). Of note, CMT2 has been previously shown to be linked with DNA methylation natural variation at many TEs^47^, and we found that these included *ATCOPIA12D*. At *AT2TE45020* (*ATCOPIA12B*), a bimodal distribution of DNA methylation was observed for the CG context in particular, most of the ecotypes being unmethylated (**Fig.EV5C**). GWAS for this TE again showed an association with a region encompassing the TE itself (**Fig.EV5D**). The observation of *cis*-peaks for both GWAS suggests that the epigenetic status could be *cis*- determined: this would support the instructive mode of H3K27me3 recruitment evidenced in Figure 3. While we cannot exclude the possibility that H3K27me3 was not determined by a particular change in the TE sequence, but rather occurred by chance in some accessions and then was maintained by robust epigenetic inheritance common in plants^48^, our insertion experiments described above that included *ATCOPIA12B* support the link between the TE sequence and the establishment of H3K27me3. Moreover, the identification of *trans*-factors further indicates that the switch is likely multifactorial.

**Figure 5.**
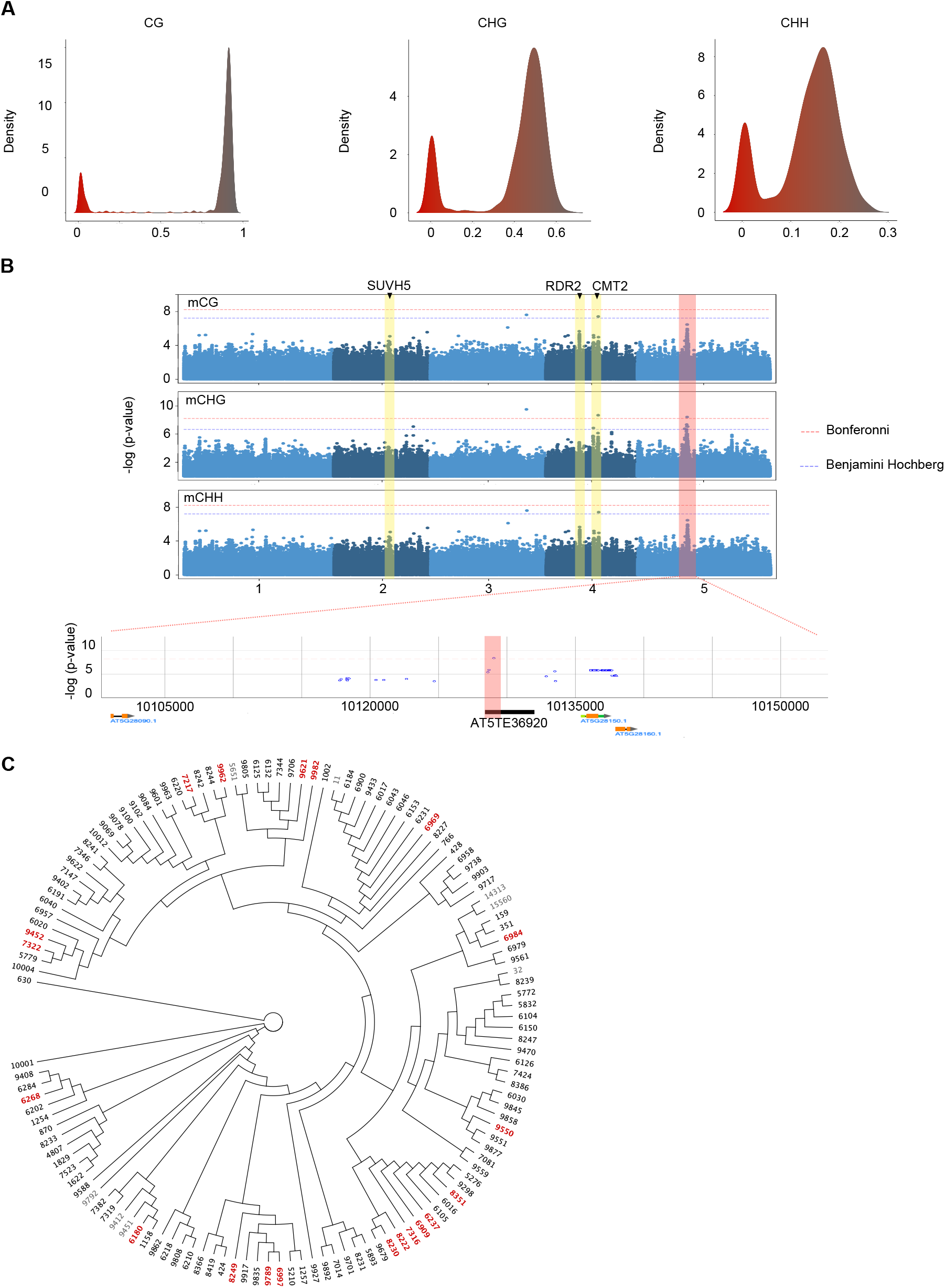
Genetic determinants of epigenetic switch (A) Plot showing DNA methylation value for an orthologous TE, *COPIA12D* (*AT5TE36920*) and the density across ecotypes in the 3 contexts, CG, CHG and CHH. **(B)** Manhattan plots show univariate GWAS mCG (**upper panel**), mCHG (**middle panel**) and mCHH (**lower panel**) for *COPIA12D* (*AT5TE36920*). Horizontal lines show genome wide significance (p = 0.05 after Bonferroni correction or after Benjamini Hochberg). Yellow boxes indicate peaks in *trans*-factors related to DNA methylation and pink boxes indicate *cis*-region of the TE. **(C)** Phylogenetic tree of AT5TE36920 sequences in 130 different ecotypes. Accessions colored in red and bold mean *COPIA12D* is devoid of DNA methylation and assumed to be marked by H3K27me3. Accessions colored in black mean *COPIA12D* is DNA methylated. Ecotypes colored in grey mean no information about DNA methylation status of *COPIA12D* (based on Kawakatsu et al., 2016)

To get more insight into the nature of possible *cis*-determinants, we extracted the sequence of *COPIA12D* in 130 assembled accessions using a previously published method^47^. We aligned them to build a phylogenic tree and observed that H3K27me3-marked sequences did not cluster together (**Fig.5C)**. This shows that there are no common haplotypes causal to the epigenetic switch to H3K27me3, and further indicates that the switch has occurred several times.

Thus, the epigenetic switches are the product of multifactorial causes leading to a change in the TE silencing mode and have likely occurred independently in nature during the species evolution.

## DISCUSSION

In higher plants, TEs marked by H3K27me3 in wild type have recently gained interest. While many H3K27me3-marked TE relics have been recently described^36^, here we report for the first time the existence of TEs that can be intact and are targeted by Polycomb instead of DNA methylation. Furthermore, our study provides multiple lines of evidence that the presence of H3K27me3 at these TEs may be linked to *cis*-determinants as shown for genes^49^. First, we observed that long H3K27me3-TEs can be found in all genomic compartments, including pericentromeric regions. Second, a transgenic approach showed that three different TE sequences display different and specific patterns of H3K27me3 upon neo-insertion. This demonstrates that PcG recruitment can be instructed by the TE copies themselves independently of position effects. Finally, and in support of this, we found that an orthologous TE copy can switch epigenetic status across accessions, independently of any epigenetic and transcriptional changes in the neighboring regions; GWAS analyses further support a link between the TE sequence and the establishment of H3K27me3. The intrinsic ability of long TE copies to recruit PRC2 raises the possibility that during the process of TE erosion, H3K27me3 nucleation regions could have been selected. Interestingly, short H3K27me3-TE relics were preferentially found nearby genes^36^. Future work should investigate whether these relics could act as epigenetic modules that can impact gene expression, as shown for DNA methylated TE remnants.

The observation that many TEs are marked by H3K27me3 raises the question of why TEs would be targeted, at times, by a plastic silencing mechanism instead of DNA methylation. When they first insert into the genome, TEs (in particular retro-elements) are DNA hypomethylated since neo-synthesized DNA (retrotranscribed) is inserted: the targeting of PcG at this stage could allow for rapid silencing of the element while DNA methylation is being established progressively. A biological role of PcG silencing is also supported by co-occurrence with H2AZ and H2Aub and the active modulation of H3K27me3 patterns by REF6/ELF6/JMJ13. As shown for H3K27me3-marked genes, this may contribute to a partially repressed state of the TE that enables a quick response to environmental or developmental cues. This may be important for organisms at certain windows of their life cycle as exemplified by TEs^50,51^ which expression is necessary during embryonic development in mammals.

Finally, inter-accessions comparisons uncovered repeated epigenetic switching between H3K27me3 and DNA methylation at a small population of TEs. Future investigations should reveal in which context such switches are triggered, although we predict that recapitulating the switch experimentally is going to be challenging. Yet, several non-mutually exclusive scenarios can already be proposed (**Fig.6A-C**). One scenario is that certain TEs have features allowing them to be targeted by Polycomb upon insertion in the genome; this is supported by our results with transgenic TEs able to recruit H3K27me3 *de novo.* Then at some point, upon TE detection by the RNAi machinery leading to onset of RdDM, and/or DNA methylation amplification, PRC2 would be antagonized (“switch 1”, **Fig.6B**). One other possible switch would entail prior loss of DNA methylation through loss of *cis* methylated cytosines and/or expression changes of *trans* modifiers, combined with *cis* molecular determinants of PcG recruitment such as PREs (“switch 2”, **Fig.6A-B**). Meeting all these requirements could take many generations. Of note, in our analyses, only a fraction of the host TEs have undergone the switch, which likely indicates that they are in a transient evolutionary state. It will be important in the future to understand how TE silencing progresses from one system to the other, because DNA methylation and H3K27me3 do not have the same properties in terms of maintenance and silencing stability. Hence, this trajectory would impact the TE potential for mobilization and domestication as regulatory modules, which could profoundly impact the integrity and expression of host genomes.

**Figure 6.**
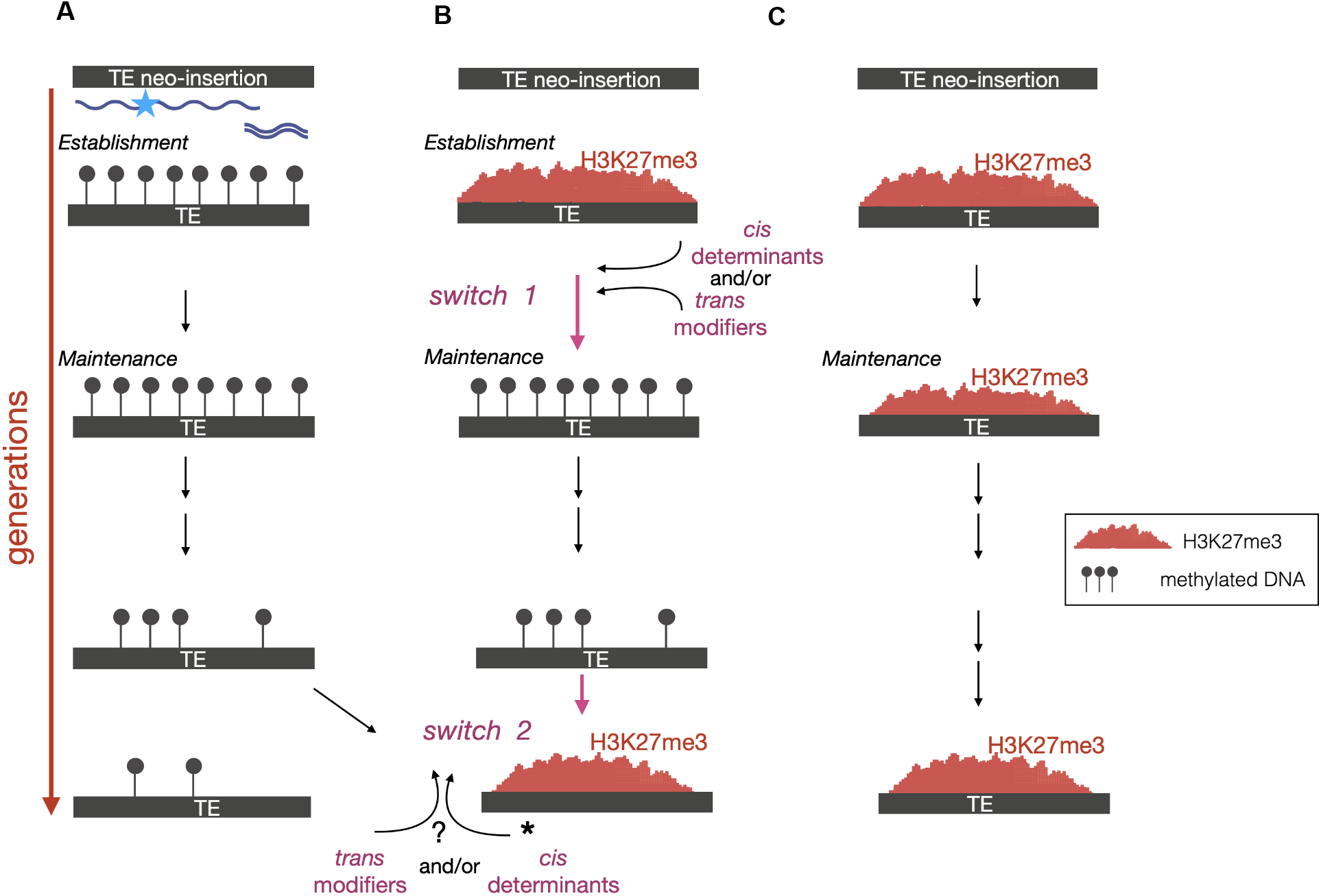
Scenario for epigenetic switch throughout TE lifetime. (A) So far, previous studies have established a model whereby TE arrive in the genome after a burst of transposition or horizontal transfer, are detected by the RNAi machinery following their transcription or by pre-existing homologous small RNAs . DNA methylation is established by small RNA-directed DNA methylation (RdDM) and maintained by a combination of RdDM and RNA-independent mechanisms. Eventually TE decay and can keep passively DNA methylation or lose it. The results obtained in this study lead to propose additional phases to this scenario of silencing progression, at least for some elements, which could comprise one or two switches “DNA methylation-H3K27me3”. **(B)** One scenario could be that PcG would be recruited first, either alone or through DRM2. This targeting could slow down the onset of DNA methylation yet DNA methylation would come in at some point, amplify through generations and become dense enough to antagonize PRC2 -this would constitute a first possible “switch 1”- and this silencing state more stable hence found more frequently. There could be another, not mutually exclusive, switch (“switch 2”) when, as part of his decaying process/by mutating, the TE loses the capacity to maintain DNA methylation and /or mutates to promote PcG recruitment. Eventually some of these H3K27me3-TEs could also be selected for structural or regulatory functions. **(C)** Some TEs may gain H3K27me3 upon insertion in the genome and keep this epigenetic status throughout their lifetime.

## MATERIAL AND METHODS

### Plant material and growth conditions

All experiments were conducted on A. *thaliana* on ½ MS plates under short light day conditions (8-h light/16-h dark photoperiod and 22°C). 4 weeks old rosettes leaves were collected in pools for the ChIP-seq transgenic approach experiment.

### Generation of transgenic lines

TE sequences were synthesized and cloned in pUC57 by Genescript. Subsequently, TEs were cloned in pCAMBIA3300 using restriction enzymes and T4 DNA ligase- based cloning strategy. Plants were transformed via *Agrobacterium tumefaciens* floral dip^52^ and T1 plants selected on appropriate herbicide for selection.

### Chromatin immunoprecipitation (ChIP) and ChIP-qPCR/sequencing analyses

ChIP experiments were conducted in WT or appropriate mutant lines using an anti-H3K27me3 antibody. DNA eluted and purified was sequenced (sequencing single-reads, 1 × 50 bp or paired-end 100bp; Illumina) of the resulting input and IP samples performed by BGI. Reads were mapped using BWA^53^ onto TAIR10 A. *thaliana.* Genomic regions significantly marked by H3K27me3 were identify using MACS2^54^ and genes or TEs overlapping these regions were obtained using bedtools^55^. Heatmap and plotprofiling were made using bedtools computeMatrix to create a score matrix and plotHeatmap to generate a graphical output of the matrix.

### Read count analyses

Reads overlapping with the SNP (between transgene and endogene) at *ATCOPIA12B, ATCOPIA13A* and *ATCOPIA21A* locus were extracted, counted and normalized by total read number using samtools view. At control regions (UBQ and FLC), reads were extracted at positions *Chr2:15,143,214* (UBQ) and *Chr5:3,178,750* (1^st^ intron FLC) respectively.

### Bisulfite-sequencing analyses

Adapter and low-quality sequences were trimmed using Trimming Galore 0.6.5. Mapping was performed on TAIR10 genome annotation using Bismark v0.22.2^56^ and Bowtie2^57^.

### GWAS analyses

GWASs were performed online on: https://gwas.gmi.oeaw.ac.at with default parameters and without transformation.

### Analyzed data sets

The public datasets were downloaded from NCBI GEO using the specified GEO accession numbers: H2A.W and H2A.Z ChIP-seq (GSE50942)^58^, BS-seq Col-0 (GSE226560)^37^, WT H2AK121ub and *atbmi1abc* H3K27me3 and H2AK121ub ChIP-seq (GSE89358)^41^, *arp6* H3K27me3 and H2AZ (GSE103361)^40^, *ref6 elf6 jmj13* H3K27me3 ChIP-seq (GSE106942)^43^ and Ecotypes related data (GSE224761)^37^.

### Phylogenetic tree and multiple alignments

130 sequences were extracted from ONT-based genome assemblies of Arabidopsis accessions^47^. The obtained sequences were aligned using MUSCLE default parameter in Unipro UGENE. Then the phylogenetic tree derived from the sequences was made by the PHYLYP Neighbor Joining Method in Unipro UGENE.

## ACKNOWLEDGEMENTS

We thank Bouché N., Toffano-Nioche C. for their help with bioinformatic analyses as part of the SPS bioinformatic work group and SPS financial support. We thank the ANRJCJC (ANR- 19-CE12-0033-01 to A.D) for funding and the Genome Biology department of Institut de Biologie Intégrative de la Cellule (I2BC) for support. We thank the services and platforms of I2BC for excellent technical support (in particular A. Almeida and F. Culot for their assistance), V. Couvreux for excellent plant care and our colleagues for discussions.

---

**Figure EV1.**
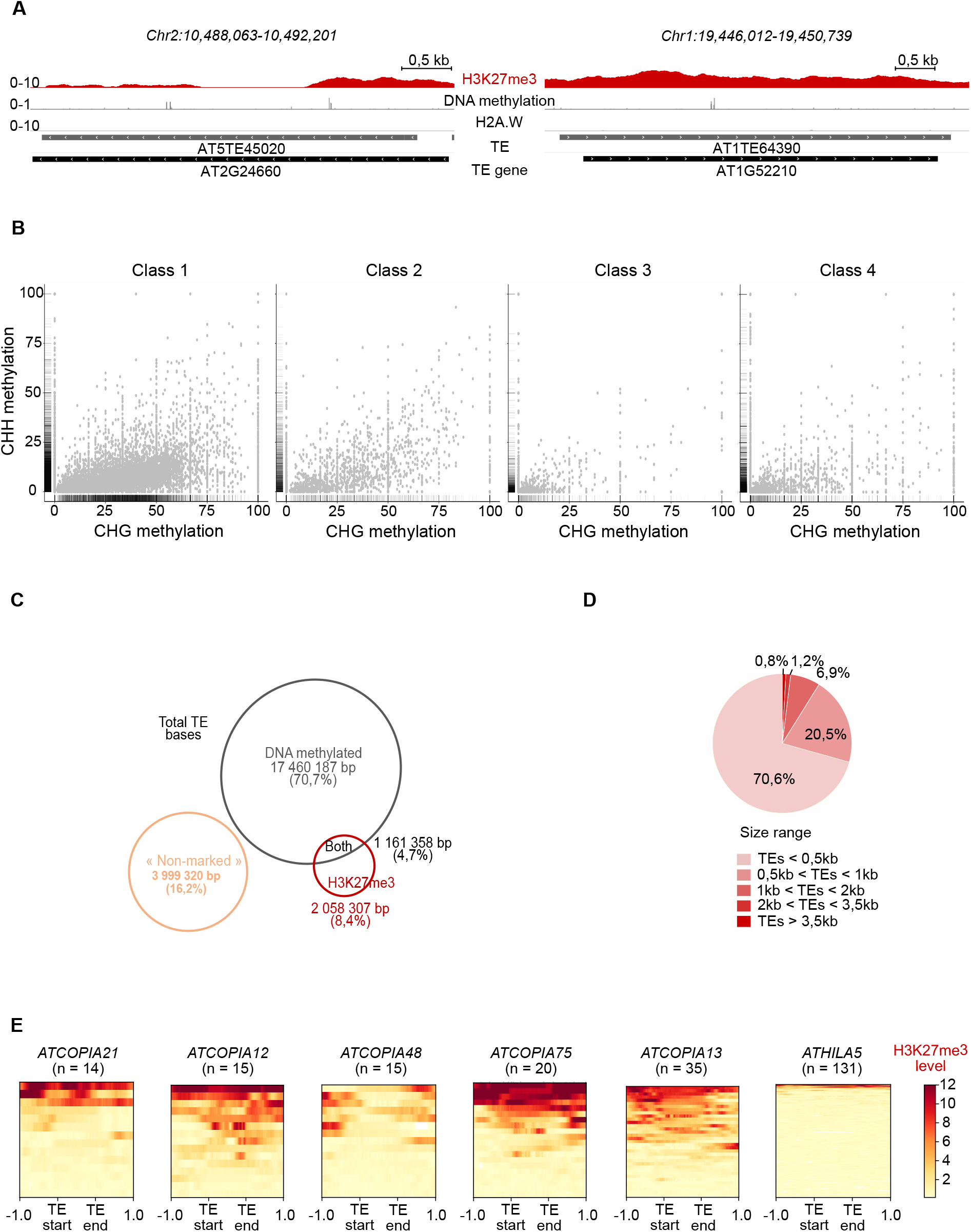
Many TE are marked by H3K27me3 in wild type Arabidopsis (A) Representative genome browser view of H3K27me3 (red), DNA methylation (grey) and constitutive heterochromatin histone variant H2A.W (grey) levels in wild-type (WT) for two TEs marked by H3K27me3 in WT. Grey and black bars represent TEs and TE-genes annotations, respectively. **(B)** Plots show CHG and CHH levels per TE in each class. **(C)** Venn diagram show the proportion of TE length for each class previously identified. **(D)** Pie chart show proportion of size range among TE marked by H3K27me3 (Class 3). **(E)** Heatmaps based on H3K27me3 levels along each TE copy within one given TE family.

**Figure EV2.**
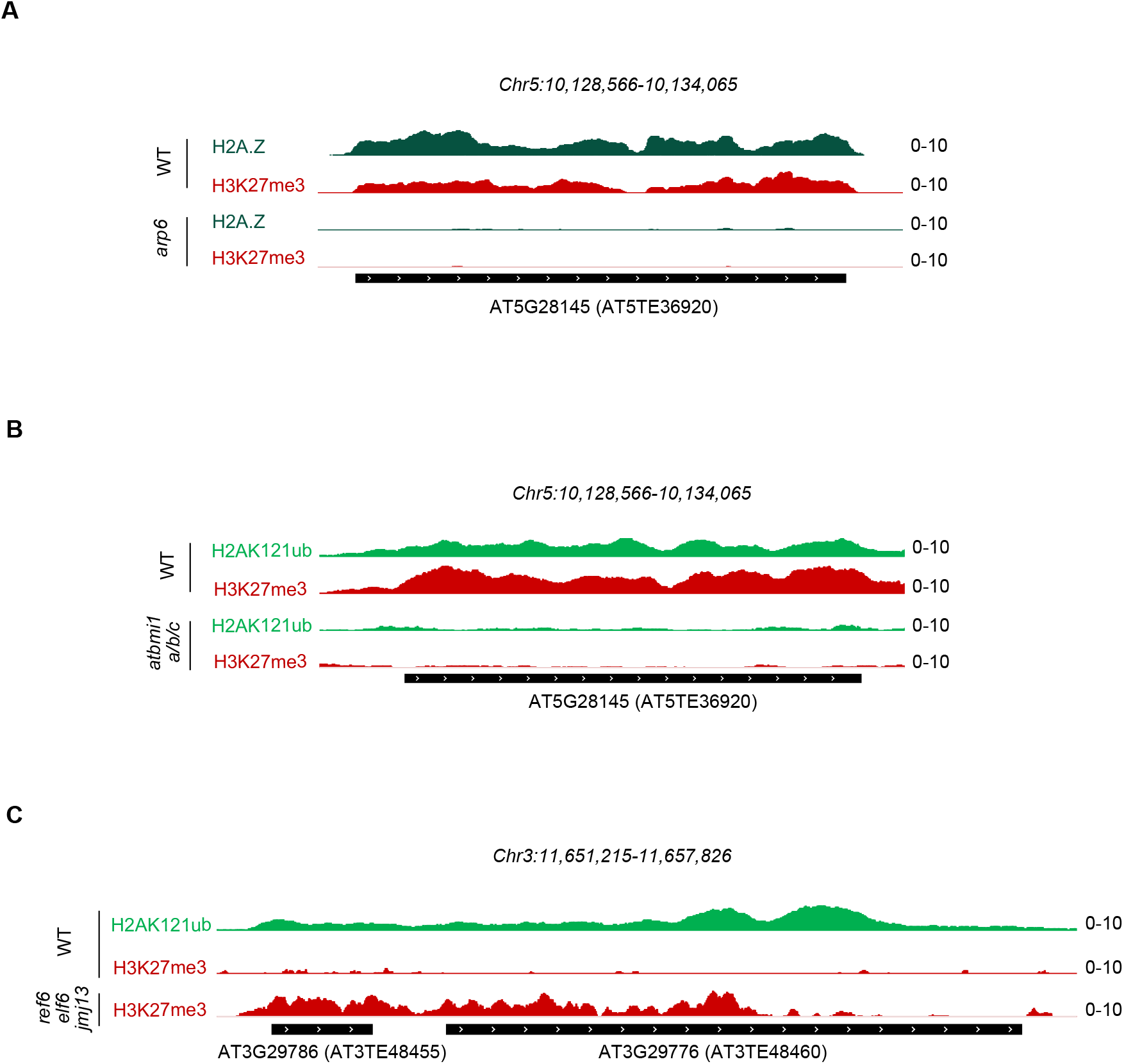
H3K27me3 at TEs can be associated with H2AUb and H2AZ and actively demethylated (A) Representative genome browser view of TE in Fig.2D with CHIP-seq data in WT and in *arp6 mutant* showing H2A.Z (dark green) and H3K27me3 (red) marks. (**B)** Representative genome browser view of TE in Fig.2E with CHIP-seqdata in WT and in *atbmi1a/b/c triple mutant* showing H2AK121Ub (light green) and H3K27me3 (red). **(C)** Representative genome browser view of TE in Fig.2F **right panel** with CHIP-seq data in WT and in *ref6 elf6 jmj13 triple mutant* showing H2AK121Ub (light green) and H3K27me3 (red).

**Figure EV3.**
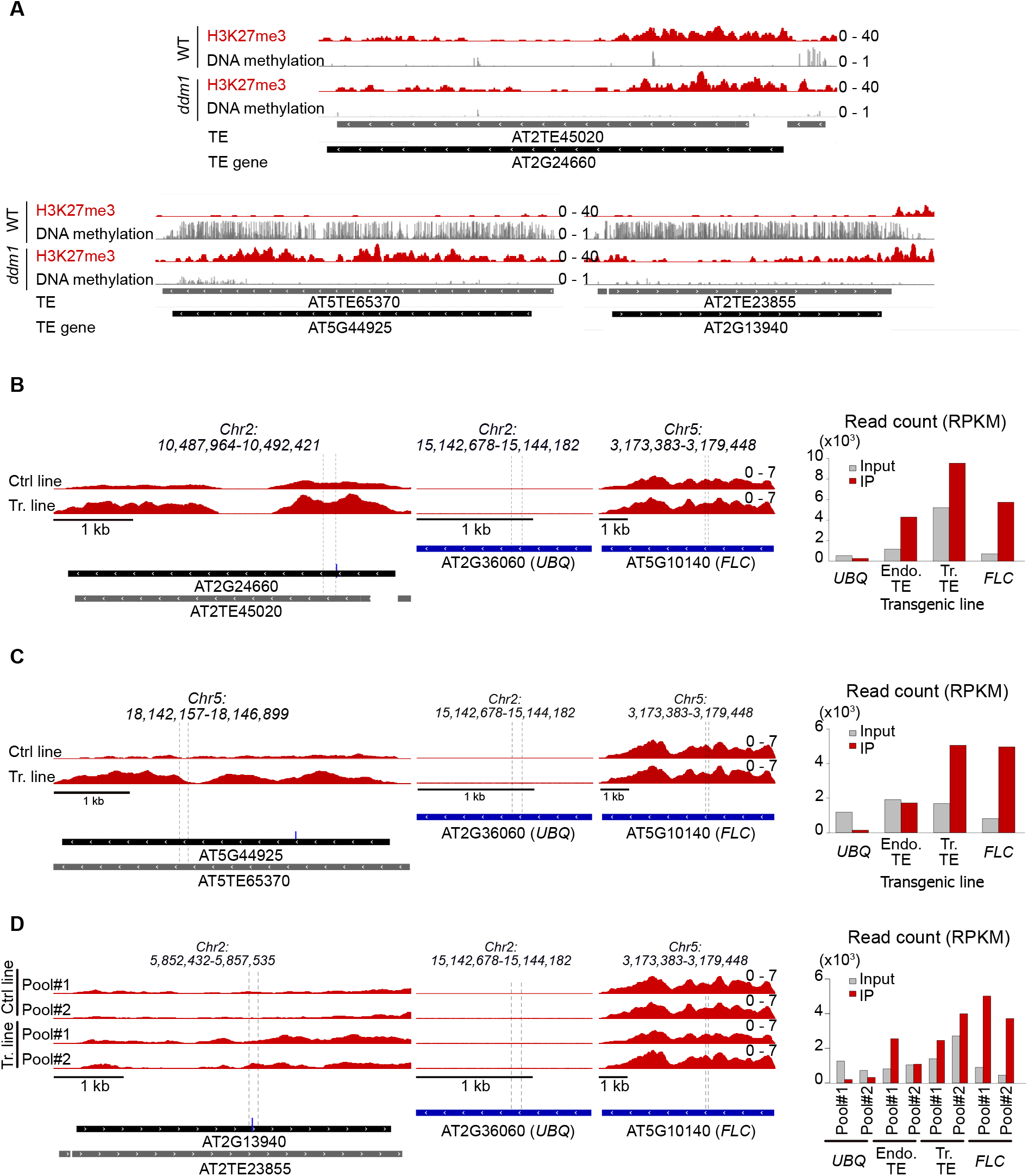
H3K27me3 can be recruited *de novo* and instructively at some transposable elements **(A)** Choice of the three individual TE sequences that were cloned into a binary vector. **(B-D)** Analyses of H37K27me3 on pools transformed with either *COPIA12B*, *COPIA21* and *COPIA13* as in Fig.3; the differences in profiles compared to Fig.3 and Fig.S2 is due to the use of another anti-H3K27me3 antibody in this experiment (Diagenode). **(B) Left panel.** Representative genome browser view of H3K27me3 average profile (IP-INPUT) at *COPIA12B* sequence (top panel) in a third independent pool of plants either transformed with unrelated transgene (Ctrl) or with the TE of interest. The H3K27me3 signal in the “Ctrl” tracks reflects the H3K27me3 status at the endogene TE. H3K27me3 signal in the “Tr” tracks reflects the H3K27me3 status at the endogene + transgene: higher signal compared to control line indicates recruitment of H3K27me3 on transgenic TEs in addition to the endogene. **Middle panel**: *UBQ* and *FLC* are shown are negative and positive controls respectively for H3K27me3 recruitment. **Right panel:** Recruitment on the transgene is by quantification of endogenous and transgenic *COPIA12B* sequences immunoprecipitated with H3K27me3 using CHIP-seq reads at the region where a SNP was introduced to discriminate transgene and endogene. Reads overlapping with the SNP at *COPIA12B* locus were extracted, counted and normalized by total read number. Recruitment on the transgene is validated by enrichment observed in IP as compared to an Input fraction. At control regions (*UBQ* and *FLC*), reads were extracted at positions *Chr5:3,178,750* (1^st^ intron) and *Chr2:15,143,214* respectively). Copy number variation between transgenic pools can be visualized by the input DNA fraction (grey bar)/ In the non-transgenic lines, variation of input reflects difference in sequencing coverage at the SNP. **(C)** Left, middle and right panels are as in (B) except for *COPIA21* sequence. **(D)** Left, middle and right panels as in (B) except for COPIA13 sequence (2 independent transgenic pools are shown as well as two control pools).

**Figure EV4.**
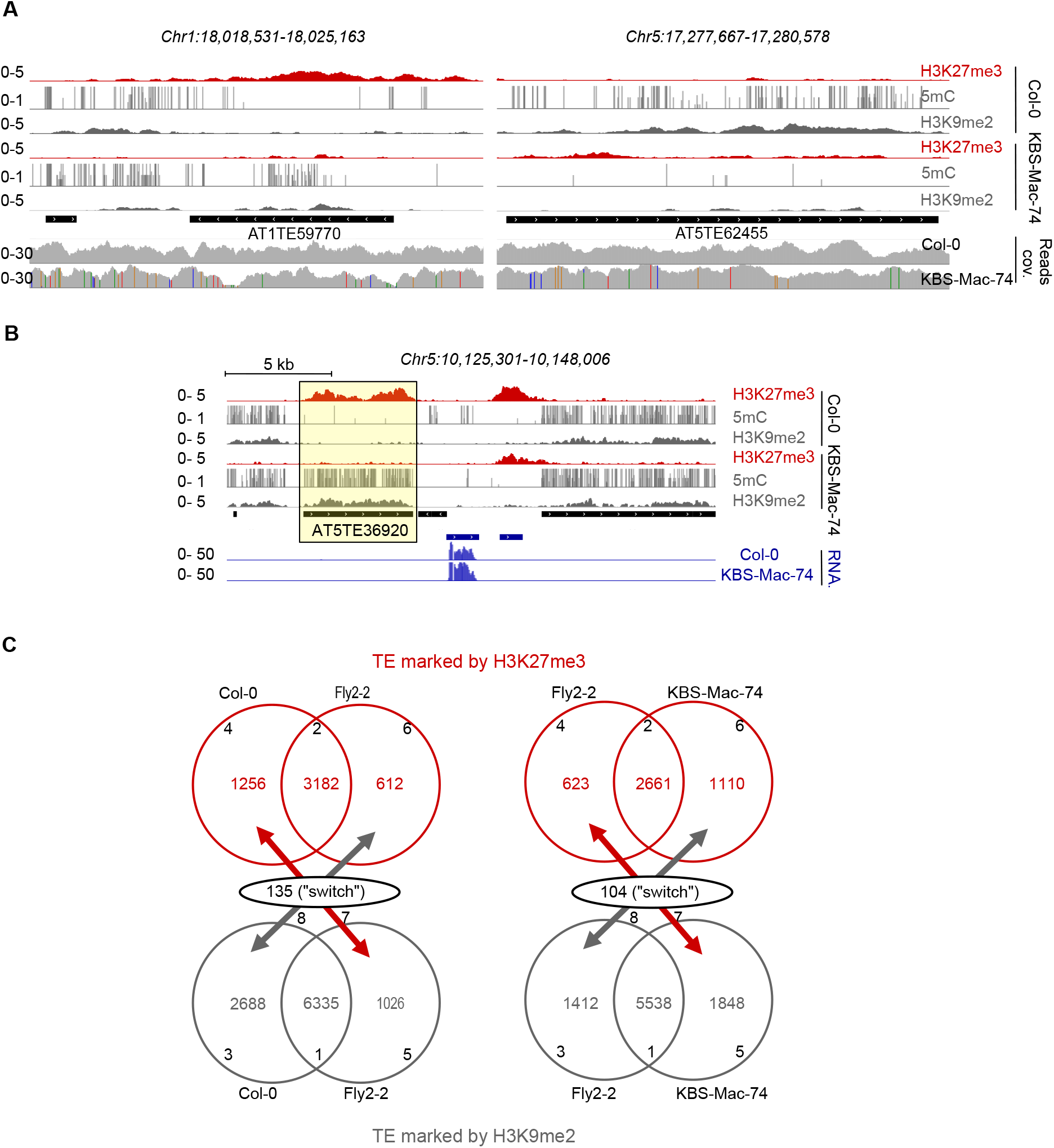
Switch between silencing epigenetic marks at TEs visualized by inter- accessions comparisons. H3K27me3 marks are in red, and DNA methylation (5mC) / H3K9me2 marks are in grey; TE annotations are shown as black bars. **(A)** Representative genome browser view of epigenetic switch at two orthologous TEs between Col-0 and KBS-Mac-74 (coverage and SNPs are shown). **(B)** Representative genome browser view of epigenetic switch at an orthologuous TE (*AT5TE36920)* with nearby regions shown, in Col-0 and KBS-Mac-74 accessions (Blue bars : Transcripts level). **(C)** Venn diagram of TEs marked by H3K27me3 (red circles), or by H3K9me2 (grey circles), in two different ecotypes, Col-0/Fly-2 (left panel) and Fly-2/KBS-Mac-74 (right panel). Intersection between both Venn diagrams show TEs that switch from H3K27me3 in one ecotype to H3K9me2 in the other.

**Figure EV5.**
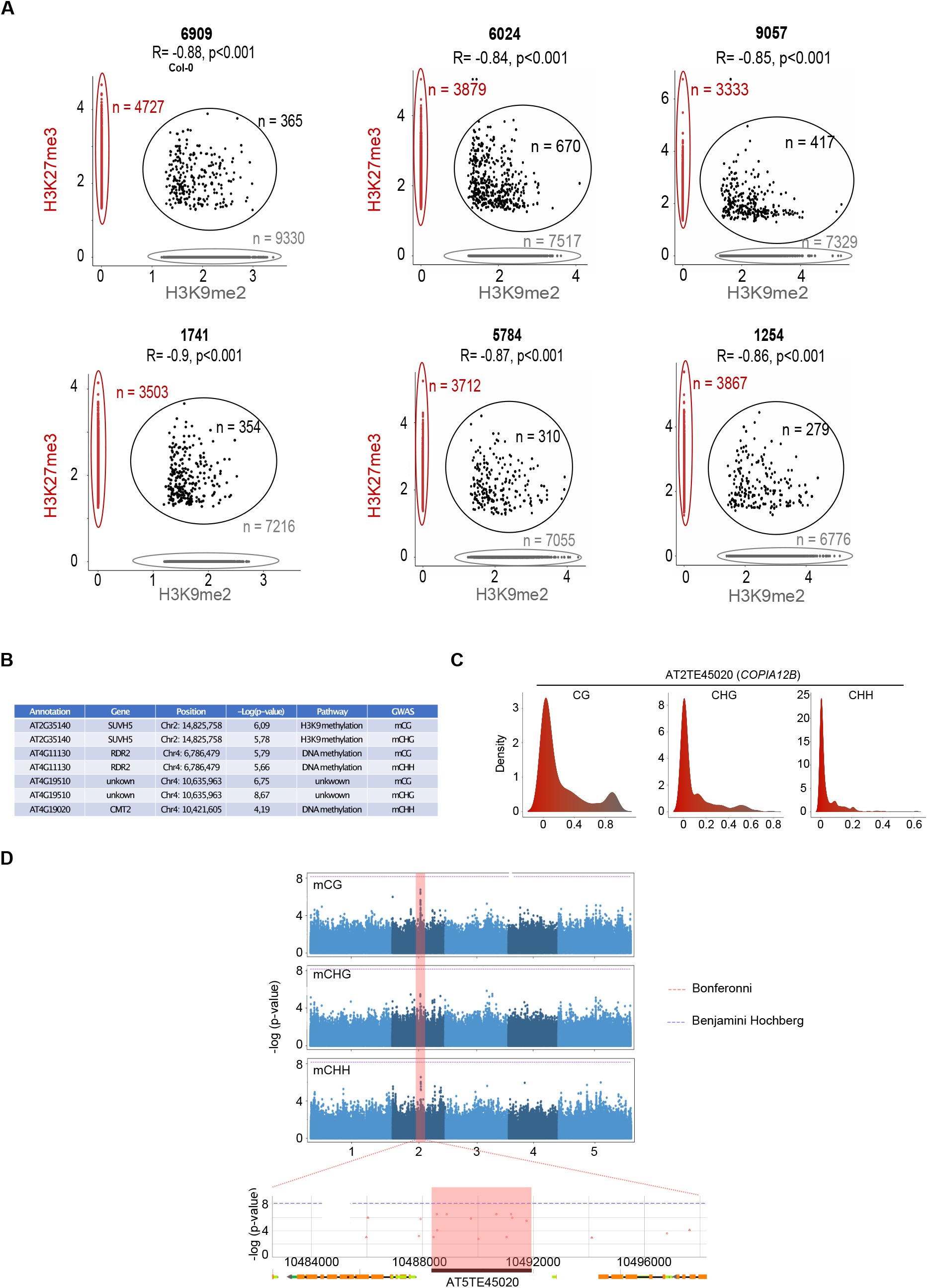
Genetic determinants of epigenetic switch (A) Plots showing correlation between H3K27me3 and H3K9me2 values for each TEs in 6909 (Col-0), 6024 (Fly2-2), 9057 (Vinsla), 1741 (KBS-Mac-74), 5784 (Ty-1), 1254 (Tos-82-387). Red indicates TE marked by H3K27me3, grey indicates DNA methylated TE and black indicates TE marked by both H3K27me3 and DNA methylation. **(B)** Table showing *trans-*peaks found in GWAS of *AT5TE39620*. **(C)** Plot showing DNA methylation value for *COPIA12B* (*AT2TE45020*) and the density across ecotypes in the 3 contexts, CG, CHG and CHH. **(D)** Manhattan plots show univariate GWAS mCG (**upper panel**), mCHG (**middle panel**) and mCHH (**lower panel**) for 2 TEs, **left panel**: AT2TE45020 and **right panel**: AT5TE36920. Horizontal lines show genome wide significance (p = 0.05 after Bonferroni correction or after Benjamini Hochberg). Yellow boxes indicate peaks in *trans*-factors related to DNA methylation to and pink boxes indicate *cis*-region of the TE.

## Notes

### Competing Interest Statement

The authors have declared no competing interest.

